# Evaluation of sources of variability in a TEER (trans-epithelial/endothelial electrical resistance) system

**DOI:** 10.64898/2026.11.06.764605

**Authors:** Robert Gutierrez, Steven Lund, Vytas Reipa, Elijah J. Petersen

**Affiliations:** Biosystems and Biomaterials Division, National Institute of Standards and Technology, Gaithersburg, MD 20899, USA; Statistical Engineering Division, National Institute of Standards and Technology, Gaithersburg, MD 20899, USA

**Keywords:** New approach methods, validation, qualification, *in vitro*

## Abstract

Trans-epithelial/endothelial electrical resistance (TEER) measurements are a key nondestructive measurement of the barrier integrity of cells grown on inserts. To support the use of TEER measurements in bioassays, we have evaluated the sources of variability in a case study involving a chamber-based TEER system to assess a gut co-culture model. Data from 23 separate experiments, each lasting 22 d, were evaluated. Multiple lines of evidence suggested minimal variability within a run, a period during which several plates were analyzed sequentially. In contrast, run-to-run variability was substantially larger, even when two runs were performed several hours apart on the same day. Bayesian analysis of both KCl and blank insert data revealed that run-to-run variability was among the largest sources of variability, with relative uncertainty values of 3 % to 6 % of the mean value. Correlational analysis investigated whether deviations of KCl or blank insert TEER values within a run compared to long-term median values were correlated with deviations of the biculture gut model values within that same run from their long-term median values. Unexpectedly, the KCl values were not correlated, while the blank insert values had a weak correlation. We also evaluated the information gained from different control measurements. The findings from this study can influence how statistical analyses are performed on TEER results and can inform the design of robust TEER methods. For example, the approaches described here can be used to characterize variability within and among runs in TEER measurements for other methods.

## Introduction

There is widespread interest in advancing cell-based methods due to their potential to better recapitulate human biology than traditional methods and to reduce reliance on animal testing (Food and Drug Administration 2025; National Institutes of Health 2025; Reardon 2025; Herron et al. 2025; EPA 2026; Kwon 2026; Burki 2026). To form a cell barrier that can be used to represent, for example, ocular, dermal, intestinal, and respiratory tissues, it is necessary to grow cells on inserts. This enables different media on the apical and basolateral sides of the insert. For example, inhalation-related methods may have air on the apical side and cell medium on the basolateral side (Polk et al. 2016; Secondo et al. 2017; Lacroix et al. 2018; Zavala et al. 2018; Leibrock et al. 2020; Petersen et al. 2021; Haber et al. 2024; Mallek et al. 2024).

Trans-epithelial/endothelial electrical resistance (TEER) measurements assess the barrier integrity of cells grown on inserts by measuring their resistance to an electrical current (Srinivasan et al. 2015; Karakocak et al. 2023; Sharma et al. 2025). Higher resistance corresponds to greater barrier integrity. There are several types of probes for TEER measurements, with different geometries such as prongs or chamber systems (Srinivasan et al. 2015; Sharma et al. 2025). These measurements can also be used in microphysiological systems (Malik et al. 2025; Torres-Castro et al. 2026). Importantly, these measurements are nondestructive. Some Organization for Economic Cooperation and Development (OECD) test guidelines using 3D constructs specify a minimum TEER value to ensure the construct integrity (e.g., (OECD 2021, 2025)).

TEER measurements can be used in a range of contexts: 1) test method specifications, 2) control measurements for the biomanufacturing of a biological sample on an insert, and 3) statistical comparisons among measurements taken at the same or different time points. There are numerous factors that could complicate these comparisons, including but not limited to changes in the sensitivity of the probe such as from degradation of the AgCl electrode, changes in the temperature of the medium used during the TEER measurement, and intentional or inadvertent changes to the biological samples that influence their barrier integrity.

Here, we characterize potential sources of uncertainty in TEER measurements of a biculture gut tract model with a chamber probe system as a case study (Figure 1). Control measurements for the different components of the TEER system were taken over nearly 2 years. We investigated the potential to correct instrument drift for measurements performed hours or days apart. This evaluation is intended to complement a recently published study on reporting guidelines for TEER measurements (Sharma et al. 2025). The reporting guidelines clarify what to report in a TEER study, while this study describes strategies for characterizing and controlling different sources of variability.

**Figure 1:**
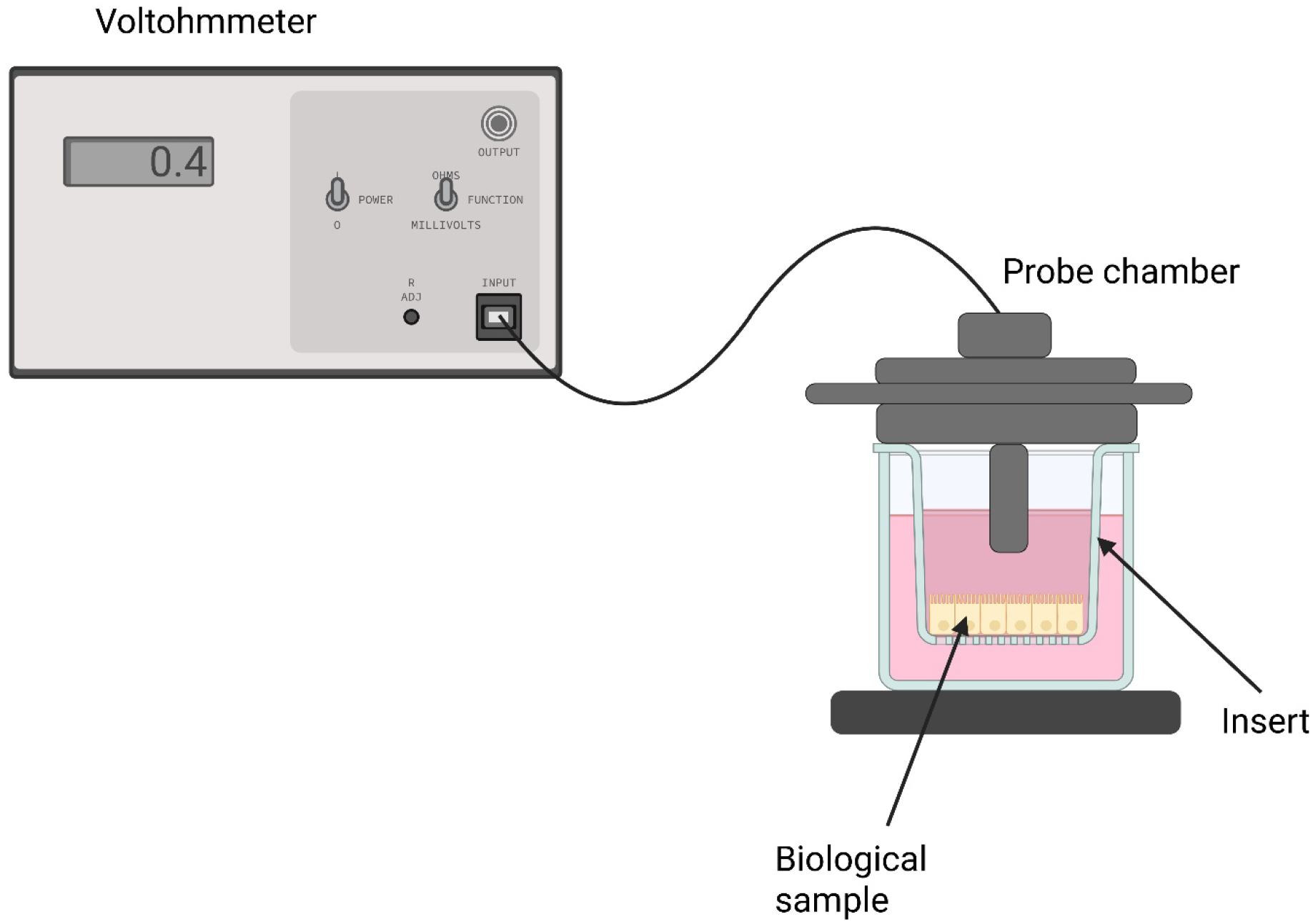
Schematic of the TEER instrument using a chamber probe. Produced using Biorender.com.

## Method

In this manuscript, the term “experiment” refers to a 22 d duration when multiple plates with different gut models were grown and periodically analyzed using TEER. The term “run” refers to the period when several plates are analyzed consecutively using TEER. After each run, the TEER probe was washed using the procedure described below. Typically, only one run was performed each day, but in the first four experiments conducted, a second run was performed 1 h (Experiment 1) or 4 h (Experiments 2, 3, and 4) after the conclusion of the first run. During each run, KCl solutions and blank inserts (i.e., inserts without cells) were analyzed as control measurements, except that KCl solutions were measured only at 22 d in Experiment 5. KCl measurements from Experiment 1 were not used because no original record was available. The original intention for the KCl solution measurements was to correct for run-to-run variability in the instrument sensitivity, while the purpose of the blank inserts was for background correction of the gut tract model results.

### Cell Culture

Caco-2 (human colorectal adenocarcinoma) cells, HT29 (human colorectal adenocarcinoma) cells, and Raji B (human B lymphocyte) cells were obtained from ATCC.

Caco-2 (passage 5 to 30), HT29 (passage 5 to 30), and Raji B (passage 5 to 30) cells were separately cultured in complete medium, consisting of Dulbecco’s modified Eagle medium (DMEM) high glucose (Thermo Fisher Scientific) with volume fractions of 20 % heat inactivated Fetal Bovine Serum (HI-FBS) (Thermo Fisher Scientific), 1 % non-essential amino acids (NEEA) (Thermo Fisher Scientific), and 1 % penicillin and streptomycin (Thermo Fisher Scientific) using a tissue culture flask (Corning). Cells were maintained in a 37 °C incubator with 5 % CO₂. Cells were sub-cultured twice a week at 90 % confluence by detaching with trypsin-EDTA (volume fraction 0.25 % Trypsin and 2.21 mmol/L EDTA; Corning) solution.

### Culture Model Assembly

Three different gut models (monoculture, biculture, and triculture) were grown in parallel, largely following previously developed protocols (Araújo and Sarmento 2013; Vincentini et al. 2022). They were typically grown on 3 µm pore size transparent PET membrane inserts (Corning) placed in 12-well plates (reference number 353181; 0.9 cm^2^; Falcon); measurements in experiments 15 and 16 and some measurements in experiment 14 were made using a different insert: 3 µm pore size PET membrane inserts (reference number 3462; 1.12 cm^2^; Corning). Monoculture models consisted of only Caco-2 cells by seeding 250 000 cells/insert on the apical side, while the biculture and triculture had Caco-2 and HT29 cells added at a 9:1 ratio [Caco-2 (225 000 cells/insert) and HT29 (25 000 cells/insert)] to the apical side. Additionally, the triculture models had 500000 Raji B cells added to the basolateral side of the inserts on days 14 and 16.

All inserts were filled with 0.5 mL of DMEM complete culture medium (DMEM high glucose with volume fractions 10 % HI-FBS, 1 % NEEA, and 1 % penicillin and streptomycin) on the apical side, while the basolateral side of the insert also had 1.5 mL of DMEM complete culture medium. The inserts were cultured for 22 d with the medium changed three times per week. For the triculture model, the Raji B cells were introduced on day 14 using a culture medium that was a 1:1 mixture of DMEM complete culture medium and Roswell Park Memorial Institute (RPMI) complete culture medium (RPMI-1640 medium (Thermo Fisher Scientific) containing by volume fraction 10 % HI-FBS, 1 % NEEA and 1 % penicillin and streptomycin). On days 14 and 16, the Raji B cells were removed from the basolateral side of the inserts, and a fresh batch of Raji B cells was added to the basolateral compartment. On day 17, the Raji B cells were removed and replaced with DMEM complete culture media; thereafter, only DMEM complete culture medium was used. The purpose of this change, compared to previous methods in which Raji B cells were added three times, was to avoid cell treatment on weekends. After each run, the inserts were stored in DMEM complete culture medium.

### Barrier Integrity

An EndOhm-12G chamber electrode device (World Precision Instruments) and an EVOM3 voltohmmeter (World Precision Instruments) were used to make periodic TEER measurements on days 14, 17, 20, 21, and 22; measurements were also made in experiments 1 through 10 after 3 d and 7 d. On these days, the cells were analyzed by TEER prior to media exchange. The samples were moved out of the incubator and placed in a biosafety cabinet for about 30 min to 45 min to allow the cells and medium to equilibrate with the temperature in the biosafety cabinet. This time period was determined to be sufficient for the samples and media to equilibrate during preliminary experiments. Blank inserts without cells were used as a control measurement. Three repeated measurements were taken per insert and biological sample. All insert and biological sample measurements were made using DMEM complete culture medium in the chamber. The medium was not replaced during a run. To minimize any potential contribution of heterogeneity within the insert to measurement variability, the inserts were placed in a consistent alignment with the chamber for each measurement.

### Equipment

The EVOM3 voltohmmeter was calibrated using a TEER control test resistor module (MedTec) that included 9 control resistance measurements with nominal values of 10 Ω, 50 Ω, 100 Ω, 250 Ω, 500 Ω, 1 000 Ω, 2 500 Ω, 5 000 Ω, and 10 000 Ω; actual values were within 0.7 % of the nominal values. At the start of many runs, the resistor module was used to ensure the EVOM3 accurately measured resistance. Given the consistency of the results, we changed from testing before each run to periodic testing. Starting in experiment 3, the temperature of the biosafety cabinet was recorded using one or two different thermometers (Cole-Parmer Traceable or Fisherbrand digital thermometer).

The EndOhm-12G TEER chamber electrode (World Precision Instruments) was used to take measurements. The chamber lid was positioned at a specific height using a 4 mm spacer disk (World Precision Instruments) and locked in place. After each run, the chamber was sterilized using 70 % isopropyl alcohol and rinsed with de-ionized water. The plates were analyzed in the following order: monoculture, biculture, and then triculture.

The cleaning and washing procedures recommended by the manufacturer were followed after each run: the EndOhm-12G electrodes were washed using distilled water for about 1 min. The electrodes were then wiped with a Kimwipe (lint-free). The electrodes were soaked in 1 % mass volume fraction Tergazyme solution (prepared from powder; Alconox) for approximately 5 min to reduce protein contamination. De-ionized water was used to rinse the electrodes three times before wiping them down with Kimwipes, allowing them to dry, and storing them in the dark. Furthermore, the electrodes were rechlorided approximately every two weeks or more often on an ad hoc basis. The electrodes were submerged in a 5 % volume-fraction solution of sodium hypochlorite (bleach) for approximately 10 min, rinsed with de-ionized water, dried with Kimwipes, and stored dry.

Measurements were also performed after repolarizing the silver chloride layer on the electrode by anodic polarization in 1 mol/L HCl solution to assess if repolarization could reduce the run-to-run variability. Following rinsing in deionized water, the electrode was immersed in a beaker containing 1 mol/L HCl. and a 2 mA/cm^2^ current was applied using M263 Galvanostat (EG&G) for 5 min. A Pt mesh was used as a cathode. Then, the electrode was left to equilibrate in deionized water for 8 h.

### Potassium chloride (KCl)

A 160 mmol/L KCl stock solution was prepared by adding 1.193 g of KCl to 100 mL of deionized water and mixing thoroughly. This stock solution was used to prepare 80 mmol/L, 40 mmol/L, 20 mmol/L, 10 mmol/L, 5 mmol/L, 2.5 mmol/L, and 1.25 mmol/L KCl solutions by 1:1 serial dilutions in deionized water. Potassium chloride solutions were stored at room temperature. A total of eight different lots of KCl were prepared separately during this study. Lots 4 through 8 were prepared independently on the same day to directly assess lot-to-lot variability in solution preparation.

### Modeling KCl results using Kohlrausch’s law

Conductometric measurements of five KCl solutions from 20 mmol/L to 1.25 mmol/L were used to verify the system’s performance across a resistance range that covered the expected values of the gut models. Since KCl is a strong electrolyte, the solution conductance is expected to follow Kohlrausch’s law: the solution’s conductance (inverse of resistance) is the sum of the conductance of the individual ions Cl^-^ and K^+^ (Bockris and Reddy 1970). Electrolyte conductance plotted against the square root of the molar concentration should be linear (equation 1)

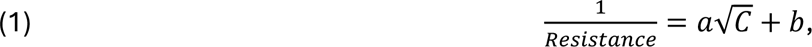

where *C* is the molar concentration and *a* and *b* are constants.

### Within- and among-run instrument drift evaluation

Experiments were performed to assess whether storing the electrode in different types of test media would impact the measured resistance values within and among runs. Four media were evaluated: DMEM, DMEM complete culture medium, DMEM complete culture medium with a monoculture cell insert, and DMEM complete culture medium with a biculture cell insert that contains mucus-producing cells. The purpose of adding monoculture or biculture inserts was to test if the presence of a cellular construct could impact the chamber electrode during a run. The measurement procedure is diagrammed in Figure 2. Initial measurements of 1.25 mmol/L and 2.5 mmol/L KCl solutions were performed, then the chamber was rinsed, and different types of media were added for 20 min, the medium and inserts removed, the chamber rinsed, and pure KCl solution conductance measurements were measured for a second time. Next, the chamber underwent the washing procedure performed after each run, and the sequence was repeated. This was performed five times in total, and the initial KCl solution conductance measurements were performed six times.

**Figure 2:**
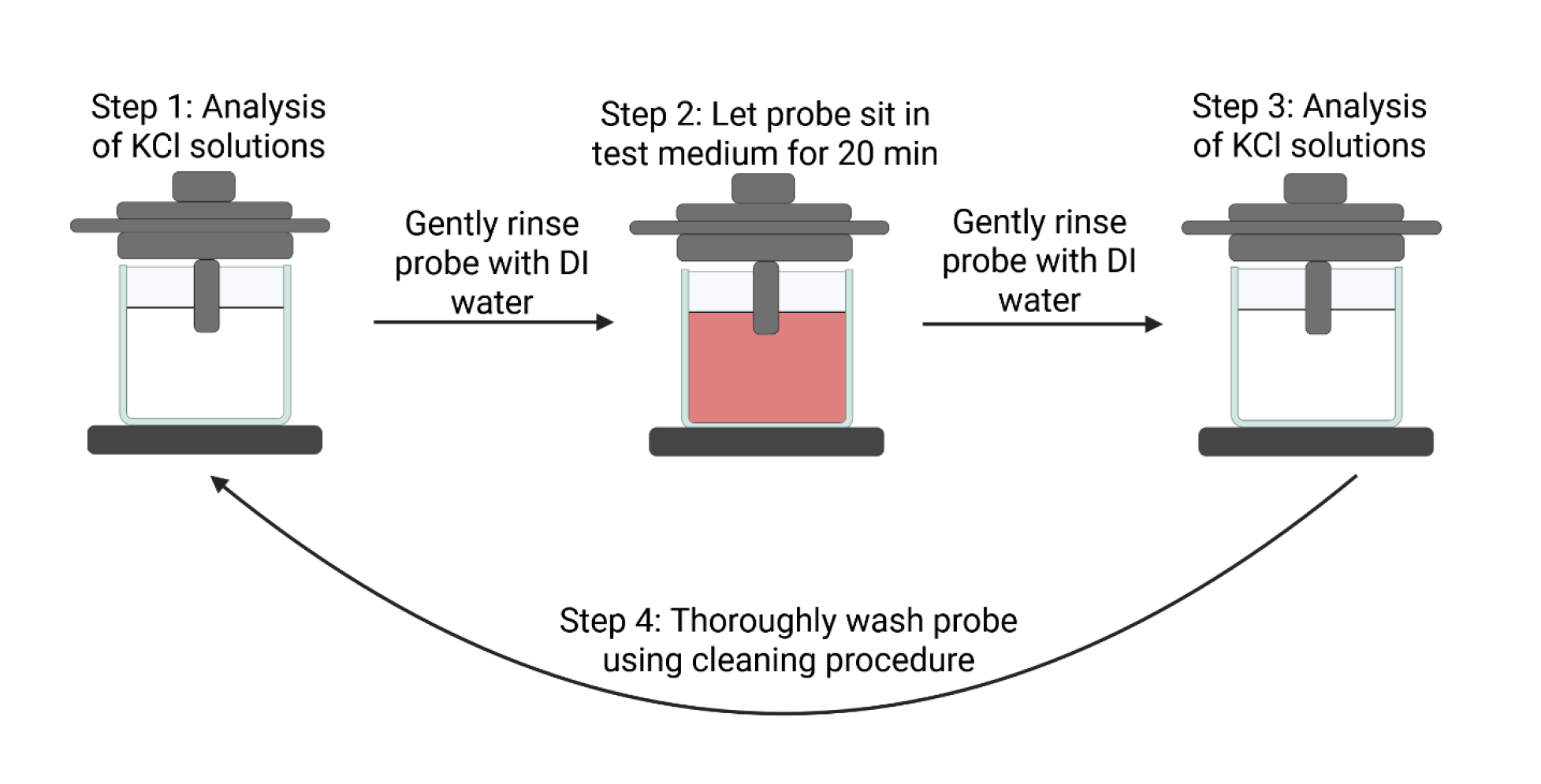
Conceptual representation of the within- and among-run instrument drift evaluation. Produced using Biorender.com.

Similar experiments were performed to assess whether electrode rechloriding or repolarization could reduce run-to-run variability compared to the simpler washing procedure. For these experiments, a CaliCell insert (World Precision Instruments) was added to a chamber with DMEM medium (without serum) and evaluated. Initial measurements of the CaliCell insert were performed, the CaliCell insert remained in the chamber for 20 min, and then the CaliCell insert was measured a second time. After each run, the chamber underwent the procedures for only washing, washing and rechloriding, or washing and repolarizing described above, and the sequence was repeated. This was performed five times in total for each treatment process, and the initial CaliCell measurements were performed six times.

Run-to-run variability for each treatment was summarized as the root-mean-square of the five log ratios between consecutive runs: the mean of each run’s initial CaliCell measurements relative to the mean of the previous run’s 20 min measurements, so each ratio spans one treatment. An increase and a decrease by the same factor count equally. Each additional treatment was compared with washing only using a one-sided F-test of whether it reduced this root-mean-square change.

### Statistical analysis

All analyses were performed in R (version 4.6.1). All analyses use data from the first run performed on a day (0 h measurements), except where noted. Readings from Experiment 11’s triculture plate at 20 days were lost due to a data transfer issue and treated as missing values. At 22 d of Experiment 13, the blank insert readings (64 Ω to 89 Ω, compared with daily medians of 18 Ω to 26 Ω in all other runs) and the KCl readings (32 % to 50 % above their lot medians, compared with at most 24 % in all other runs) were far outside their usual ranges, indicating a problem with the measurement system during that run; all measurements from that run were excluded. When plates from two different experiments were analyzed within one run, the KCl measurements were counted once in the KCl analyses. TEER values for the biological samples were calculated by subtracting the mean resistance of the blank inserts measured in the same run and multiplying by the insert area (0.9 cm^2^). Blank inserts on the triculture model plate of experiment 4 had values outside the typical range from 17 d to 22 d and were not used in this correction on those days; the inserts had likely become impacted by biomaterials during culture and had values outside of the typical range. Blank insert values were multiplied by the insert area without subtraction, as blank media subtraction was performed by the instrument. All analyses of the biological samples used blank insert-corrected values.

The relationship between conductance and KCl concentration was characterized by regressing conductance (1/resistance) on the square root of the concentration, with and without an intercept term. Individual measurements from all KCl runs in Experiments 2 through 23 were used. To estimate the typical resistance at each KCl concentration, each experiment was first summarized to its mean resistance at that concentration. These experiment-level means were then used to compute 95 % confidence intervals for the population mean and 95 % prediction intervals for a new experiment’s mean. Working from experiment-level means rather than from individual triplicate measurements avoids treating non-independent values as independent observations.

Temporal stability of the KCl resistance values was tested for each combination of eight lots and five concentrations. The aliquot-level median resistance was tested against calendar day using Kendall’s rank correlation. This test detects monotonic change without assuming a particular functional form. Forty such tests were performed, so p-values were adjusted for multiple comparisons using the Benjamini–Hochberg procedure (Benjamini and Hochberg 1995), giving a q-value for each panel. A q-value is the smallest false discovery rate (the expected fraction of false positives among the tests declared significant) at which that test would be declared significant.

A Bayesian random-effects model fit in Just Another Gibbs Sampler (JAGS, version 4.3.1) was used to estimate variability from each source in the experimental design. The model used individual resistance measurements from all eight prepared lots. Before fitting, resistance values were divided by the median for that concentration and log-transformed. With this scaling, a standard deviation of 0.04 corresponds to about 4 % variability. Variability was partitioned into three sources: lot-to-lot, aliquot-to-aliquot within a lot, and repeated-measurement noise. The model used vague priors: a normal prior on the overall mean (precision 10⁻⁶) and half-Cauchy priors on each random-effect standard deviation (scale = 1), following (Gelman 2006). The model was fit using five parallel chains with 1 000 adaptation iterations, 50 000 burn-in iterations, and 250 000 samples per chain. Convergence was checked using the Gelman-Rubin diagnostic (Brooks and Gelman 1998). Results are reported as posterior means and 95 % credible intervals. The analysis was run both with and without lot 1, which had larger repeated-measurement variability than later lots.

A separate experiment tested whether the probe’s readings change after repeated washing and exposure to different sample media as described in the “*Within- and among-run instrument drift evaluation”* subsection. Drift was monitored using reference KCl solutions at 1.25 mmol/L and 2.5 mmol/L. For each sample type and reference concentration, the per-run median resistance was tested against run number using Kendall’s rank correlation, the same test used above for the temporal stability analysis. This analysis evaluated if there was a consistent increase or decrease with an increased number of runs.

Variability in the blank insert resistance values was partitioned using the same Bayesian random-effects framework described above for the KCl resistance values. The dataset included blank insert measurements from experiments 1 through 23, with the following exclusions: a different insert product was used in experiments 15 and 16, and some plates (8 and 9) in experiment 14; experiment 13 outlier measurements; and experiment 4 triculture plate (plate 4) at the final four time points. The reference median was the overall median across all included blank insert measurements, since blank inserts are nominally constant over time and among inserts. Variability was partitioned into five sources: experiment-to-experiment, run (the same calendar date, which may span multiple experiments), plate-to-plate within an experiment, insert-to-insert within a plate, and repeated-measurement noise. Priors and MCMC settings were identical to those used for the KCl model.

Within-day variability of the biculture model was assessed using data from Experiments 1, 2, and 4, in which a second run was performed on the same day (Experiment 3 had no biculture samples due to an operator error). The second run at 22 d of Experiment 2 was excluded because it was made in a different medium. For each experiment-by-day combination with biculture measurements in both runs, the ratio of the first-run mean to the second-run mean was computed. These ratios were then summarized across experiments and days.

To assess whether run-to-run shifts were similar across sample types (e.g., the blank inserts had a similar change as the biculture samples), a run-level scaling factor was computed for each sample type as a log ratio. The blank insert scaling factor was the log of the run-level median blank resistance divided by the overall median blank resistance across the study, since blank inserts are nominally constant over time. The KCl scaling factor was the median, across all KCl measurements in the run, of the log ratio of each measurement to the overall median for its concentration. The biculture, monoculture, and triculture scaling factors were each defined analogously to KCl, with the long-run reference set to the median for that sample type on the same experimental day across all experiments. On this scale, a value of 0 corresponds to the long-run typical value, a value of −0.04 corresponds to roughly 4 % below typical, and a value of +0.04 corresponds to roughly 4 % above. Pairwise relationships between scaling factors were assessed using Pearson correlation, with significance evaluated by an F-test for the slope of the corresponding linear regression. Figure axes are relabeled in ratio units (e.g., 0.96, 1.00, 1.04) for interpretability.

Linear correlation analysis between mean blank insert or biculture TEER values and mean biosafety cabinet temperature values among runs was performed in R. When two thermometers were used or when readings were made at the beginning and conclusion of the run, their readings were averaged. Blank insert and biculture data were excluded as described above. Because biculture TEER changes with time in culture, each run’s mean biculture TEER was divided by the median of the run means for that day of culture.

## Results

### Voltohmmeter

Analysis of the nine control resistor measurements showed nearly identical results across the study. The values were consistently within 0.1 % of each other for each control resistor.

### KCl

Five KCl solution concentrations yielding resistance values spanning the range observed across the different biological samples were evaluated (Figure 3). Fitting this data with the linear model representing Kohlrausch’s law (R^2^ = 0.97; p < 2 × 10^-16^) produced estimated coefficients of *a* = (3.892 ± 0.015) × 10^-3^ mol^-1/2^·Ω^-1^ and *b*= (-4.203 ± 0.043) × 10^-3^ Ω^-1^; values indicate mean and uncertainties one standard deviation. However, the data showed curvature, which impacted the fit quality (Figure S1). The negative intercept term, *b*, could stem from additional small resistance in the measurement circuit, such as contact and/or wire resistance. Manufacturer specifications are provided for 20 mmol/L: 67 Ω ± 26 Ω (World Precision Instruments 2025), a value close to the mean value determined here (72 Ω) (Figure 3).

**Figure 3:**
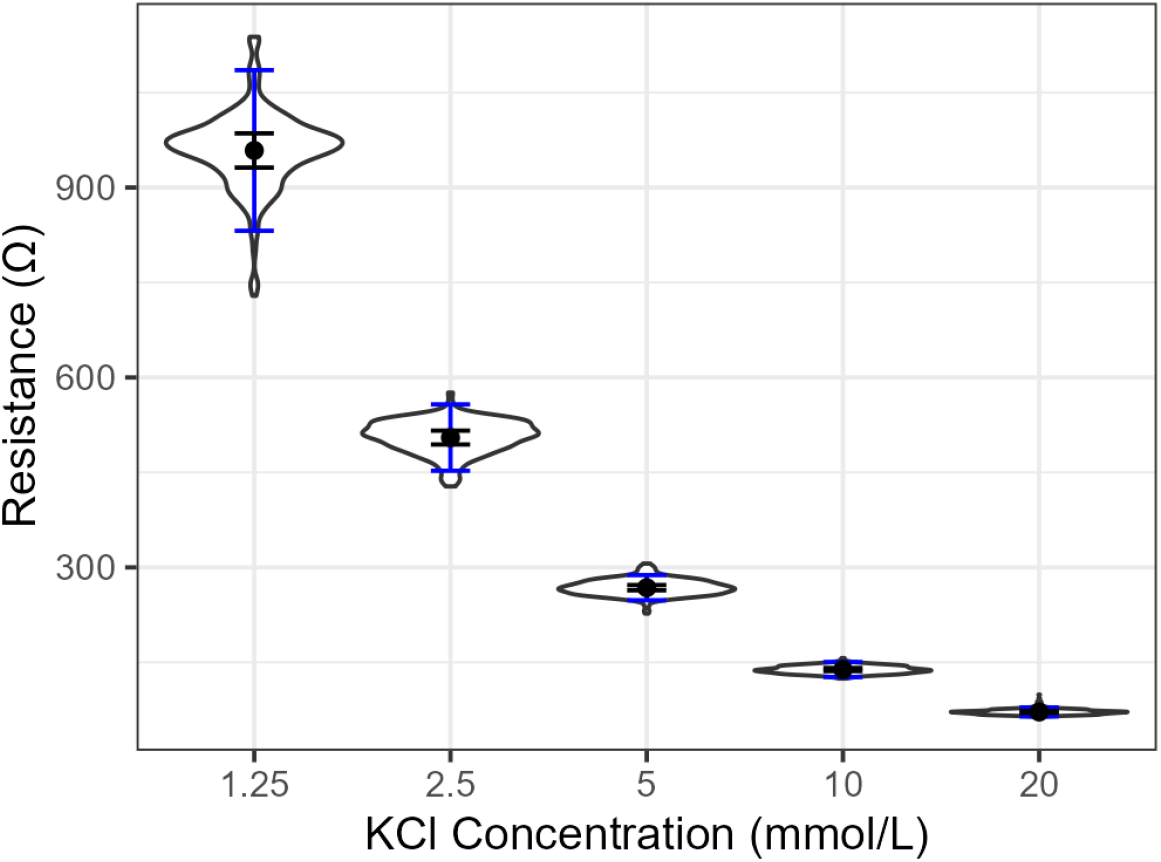
Confidence intervals and prediction intervals for the mean resistance values as a function of KCl concentration. Violin plots show the distribution of individual measurements (n=3 replicate measurements per run for data from 22 experiments; 21 at 1.25 mmol/L). When computing intervals, the data were first averaged within each experiment. Then 95 % confidence intervals for the population mean (black error bars) and 95 % prediction intervals for a new experiment mean (blue error bars) were computed across experiment means. Points show the mean of the experiment means.

This data was also evaluated to assess the stability of the resistance values across time for a particular lot (Figure S2). Unexpectedly, 28 % (11 out of 40) of Kendall correlations were significantly different from 0 (p < 0.05); after adjusting for multiple comparisons, 18 % of the panels (7 out of 40) showed a statistically significant trend at a false discovery rate of 5 %. Of the 11 panels with p < 0.05, nine had a negative slope, all at 1.25 mmol/L to 5 mmol/L, while both statistically significant trends at 20 mmol/L were positive.

Figure S3 shows the variability of the KCl resistance measurements within and among runs. Within a run, the three repeated readings of each solution varied more for lot 1 than for later lots (median coefficient of variation of 0.6 %, compared with 0.2 % to 0.4 % for lots 2 through 8), possibly because greater operator expertise increased precision for later lots (Figure S3, top). Among runs, the median of a run’s three readings was within 6 % of the median of all readings for that lot and concentration in 90 % of cases, but 24 of 704 differed by more than 10 %, the largest by 24 % (Figure S3, bottom).

Bayesian analysis was performed to assess the relative contributions of different sources of uncertainty in the resistance values for different KCl concentrations: repeated analysis of a solution aliquot during a run, run-to-run variability using the same lot at different time points, and lot-to-lot variability (Figure 4). Given the higher variability in the repeated analyses of the same aliquot within a run for lot 1 (Figure S3), this analysis was conducted with and without this lot. Including lot 1 increased the estimated variability from repeated analyses of a single aliquot by 14 % on average (ranging from a 1 % decrease to a 24 % increase among the KCl concentrations).

**Figure 4:**
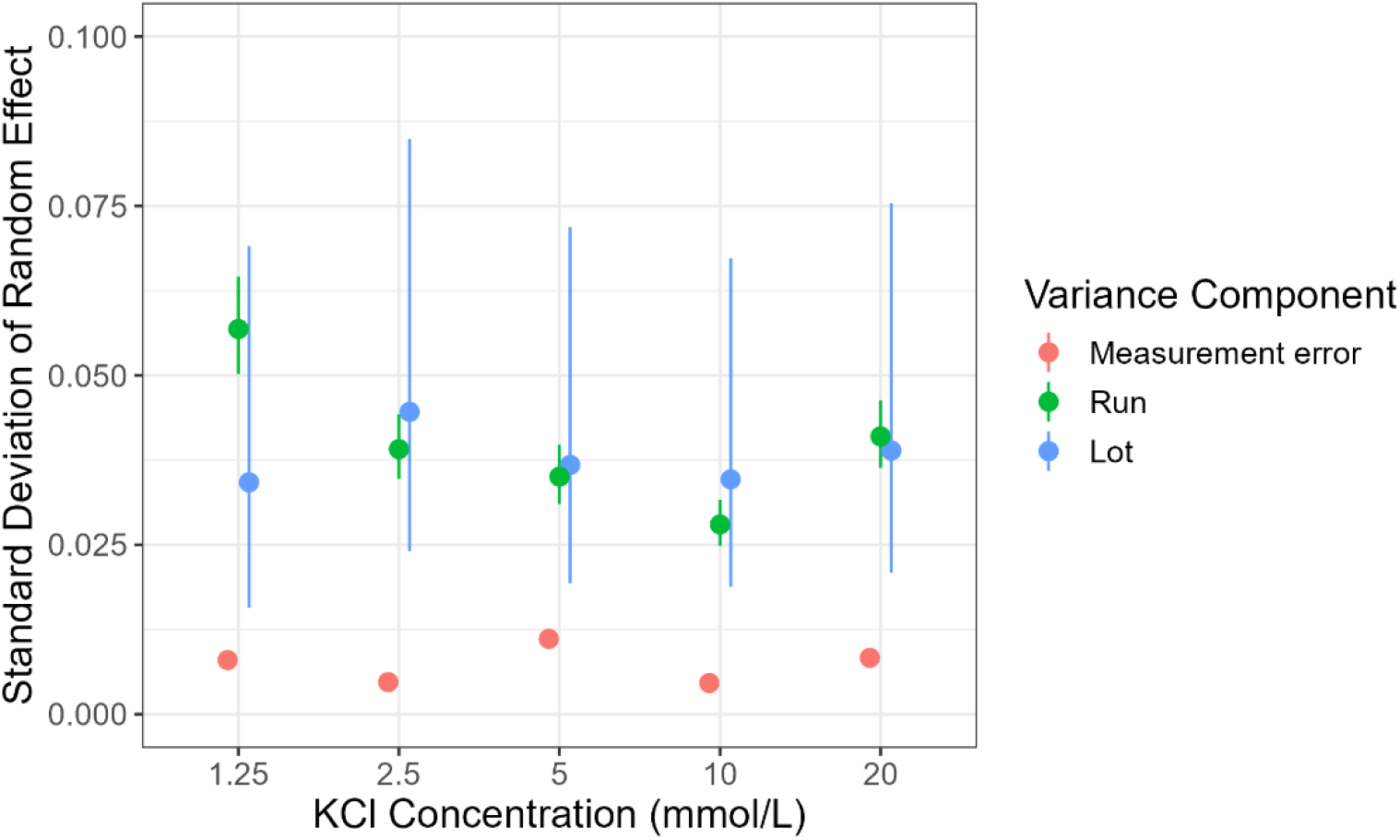
Decomposition of the variability in KCl resistance measurements (three replicate measurements of each solution in each run; 22 experiments) into three sources: lot-to-lot, run-to-run (within the same lot), and measurement error (repeated measurements of a single aliquot). The data points indicate the estimated mean standard deviation, and error bars the 95 % posterior credible interval, from a hierarchical random effects model. All analyses were performed on log-transformed data, normalized by the overall median resistance for each KCl concentration, so the standard deviation values approximate relative (proportional) variability (e.g., a value of 0.04 corresponds to a standard deviation of roughly 4 % of the overall median).

The uncertainty among runs for the same lot is largest for the lowest KCl concentration (1.25 mmol/L). This may be because this data showed a statistically significant decrease across time for lots 2, 3, and 4 (Figure S2). There is a trend of decreasing run-to-run variability from 1.25 mmol/L to 10 mmol/L with a two-fold decrease, but this trend reverses for 20 mmol/L. The contribution from the repeated measurements is fairly similar on a relative basis across the different concentrations, ranging from approximately 0.5 % to 1 % of the median value for that KCl concentration across all lots. The impact of the lot-to-lot variability was roughly an order of magnitude larger, approximately 4 %, across the KCl concentrations with overlapping 95 % posterior credible intervals.

### Drift within and among experiments

Measurements were performed to investigate the potential impact of instrument drift within and among runs using KCl solutions (Figure 5). The impact of drift within a run was assessed over 20 min, the approximate time required to take triplicate measurements of each insert in a 12-well plate. Overall, there were no clear patterns in the drift within a run. While the initial and 20 min data points often differed, there was no clear trend of increasing or decreasing values across time. In contrast, there was a clear trend of decreasing TEER values among runs for both KCl concentrations tested, 1.25 mmol/L and 2.5 mmol/L. Slopes were negative in all eight panels and statistically significant in four (Kendall, p < 0.05), ranging from 1 % to 6 % per wash. In this investigation, the probe was not rechlorided by storage in bleach (see description above), while this was performed approximately biweekly during cell measurements. While the hypothesis had been that more complex media would have a greater impact on the electrode surface condition within and among runs, there were typically not statistical differences in the slopes among the four conditions.

**Figure 5:**
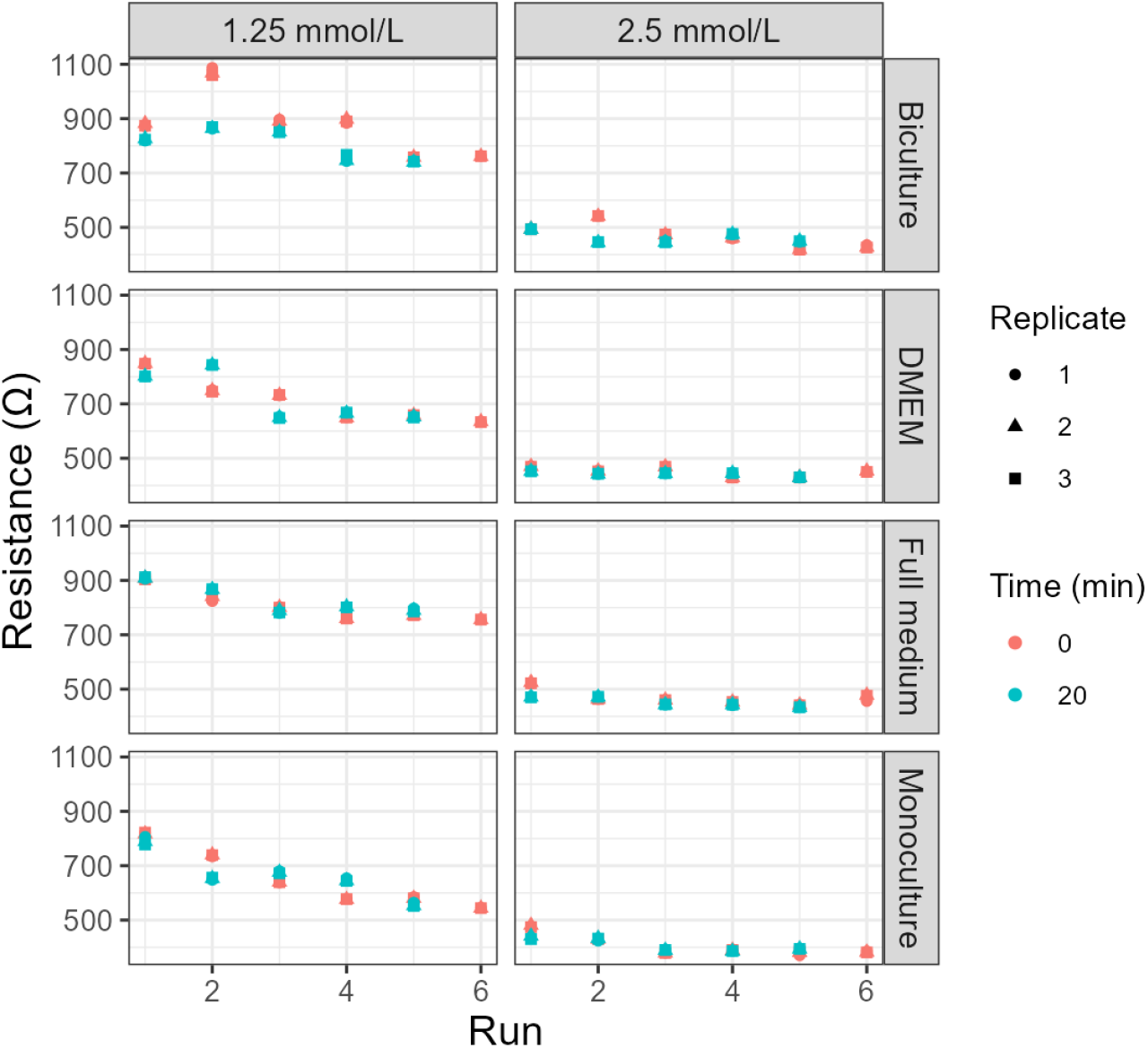
Investigation of the impact of instrument drift within and among runs. Measurements followed the scheme described in Figure 2. Triplicate measurements were made at each time point, and each is plotted individually (point shape indicates the replicate). Color indicates the time of measurement (0 min or 20 min); run 6 has only the 0 min measurement. For the monoculture condition, it was evaluated twice; the second series is shown.

Given the run-to-run variability observed in these experiments, additional experiments were performed on CaliCell inserts to investigate if rechloriding or repolarizing the electrode between runs would reduce the run-to-run variability. The drift between the initial and 20 min data points was always within 3 %. Rechloriding and repolarizing the electrodes between runs did not clearly reduce the run-to-run variability: the root-mean-square changes between consecutive runs were 6 % for washing only, 14 % with rechloriding, and 9 % with repolarizing (one-sided p = 0.95 and 0.76 for rechloriding and repolarizing, respectively, for a reduction compared to washing only). With five changes per treatment, these estimates are imprecise. Each approach (washing only, washing and rechloriding, and washing and repolarizing) yielded run-to-run differences among consecutive runs usually less than 10 % (80 % of comparisons), but 20 % of the comparisons were greater than 10 %.

### Inserts

TEER measurements were generally consistent across experiments, with an overall mean insert-blank value of 21 Ω·cm^2^ (Figure 6). Outlier results were observed in experiment 4, where the inserts were stored in complete DMEM cell culture medium during the 22 d period. In this experiment, the values of the blank inserts in only the triculture plate increased over time. A plausible explanation is that the culture medium interacted with the insert, likely clogging pores with the serum proteins, leading to higher TEER values. The blank insert values for the 22 d time point were twice as large as the values measured at 3 d and those measured in the monoculture and biculture plates at 22 d. This trend was not observed in other experiments. The affected measurements (17 d to 22 d) were excluded from Figure 6 and from the variance analysis. Outlier values were observed for the KCl and blank insert data for Experiment 13 for the 22-day time point, but the cause of this result was unclear.

**Figure 6:**
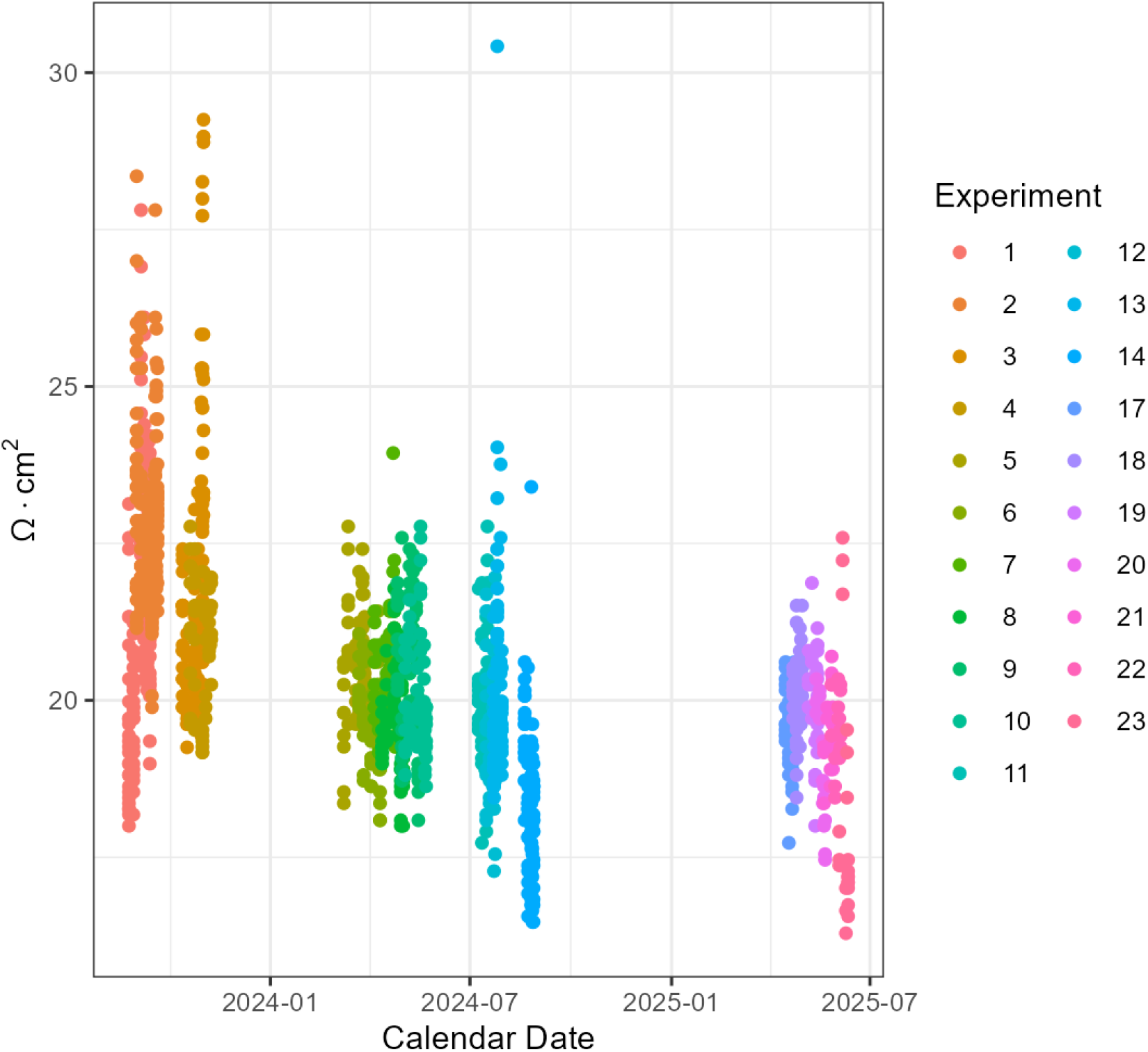
Control charting of blank insert data across time. Typically, the same inserts were used within an experiment, and different inserts were used among experiments. Data from experiments 15 and 16, and two plates from experiment 14, were excluded because these experiments involved inserts from a different manufacturer. Also excluded were plate 4 of experiment 4 at 17 d to 22 d (inserts fouled during culture), experiment 13 readings made in a different laboratory, and readings from the second run performed on the same day for experiments 1 to 4. Each data point is a single measurement of a blank insert.

The repeated measurement error of the inserts was one of the largest factors at approximately 4 % (Figure 7), a value approximately an order of magnitude larger than those for the KCl solutions. The run-to-run uncertainty within an experiment was 4 %. This is similar to the run-to-run uncertainty within the same lot for the KCl values, which spanned from approximately 3 % to 6 % even though there were sometimes statistically significant trends across time and each KCl lot, and could have been used up to 3 times longer than a single experiment; after each experiment, the blank inserts were replaced. There was also a similar uncertainty contribution from the experiment-to-experiment uncertainty, approximately 5 %, as for the KCl lot-to-lot variability, approximately 3.5 % to 4.5 %. The contributions from the plate-to-plate differences within an experiment were quite small, 1 %, as were the insert-to-insert differences within a plate, 2 %. The small plate-to-plate contribution to the insert uncertainty suggests the lack of systematic trends in the drift during runs with multiple plates. The order of blank insert measurements varied: for some experiments, all blank inserts were measured at the beginning and were on the same plate; however, for other experiments, the blank inserts were on different plates and were measured at different times during that run. Therefore, some of the contributions from plate differences could stem from instrument drift. Since the plate-to-plate effect was small, this was not investigated further.

**Figure 7:**
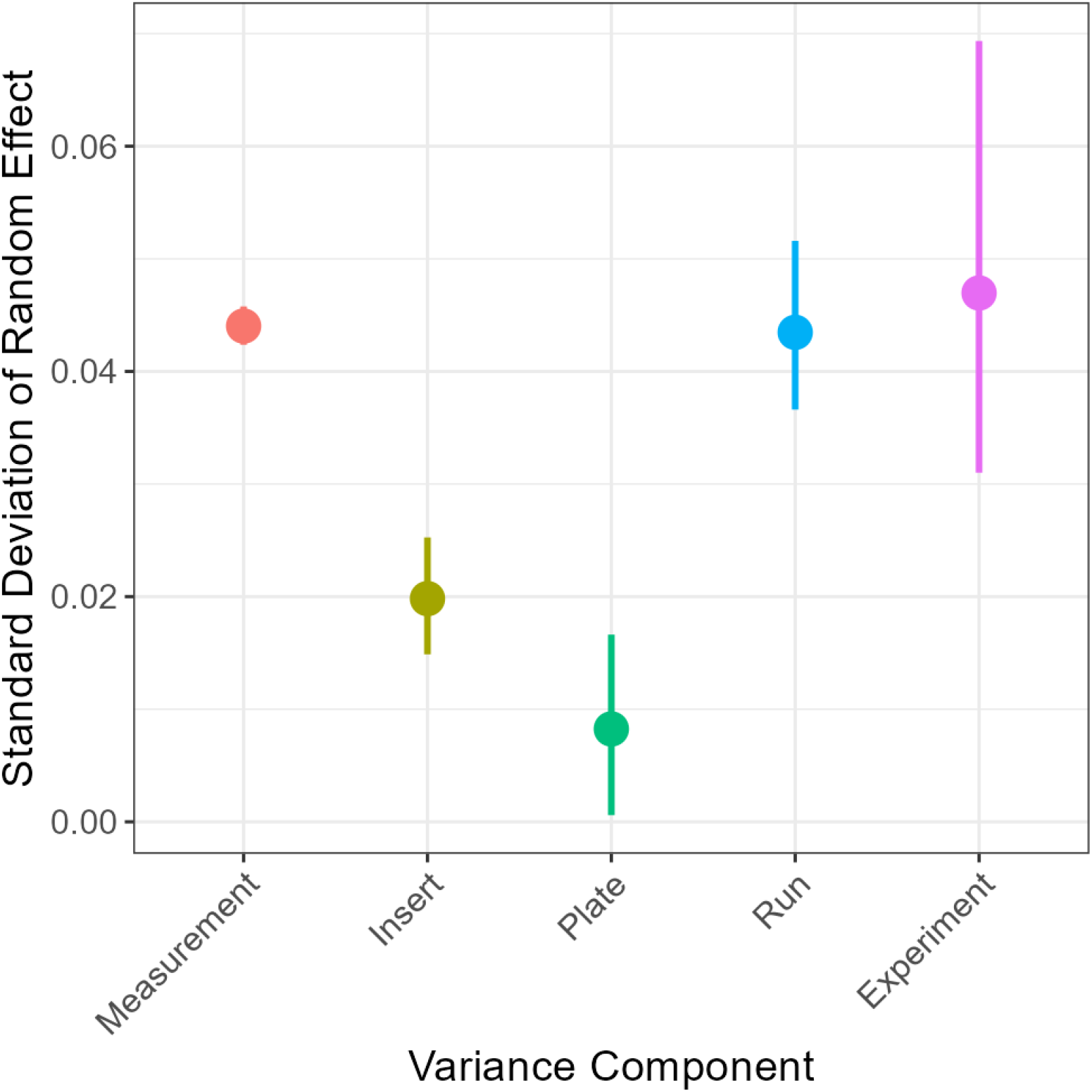
Decomposition of the variability in blank insert resistance measurements (triplicate measurements were made on each insert in most runs; in most runs of experiments 19 through 23, a single measurement was made) into five sources: experiment (variability from one full experiment, and set of inserts, to another), run (the calendar date of measurement; a run can include plates from more than one experiment), plate (variability among inserts on different plates in the same run), insert (variability between individual inserts on the same plate), and measurement (random noise among repeated measurements of the same insert). The data points indicate the estimated mean standard deviation, and the error bars the 95 % credible interval. All analyses were performed on log-transformed data normalized by the overall median resistance from all blank measurements, so the standard deviation values approximate relative (proportional) variability (e.g., a value of 0.04 corresponds to a standard deviation of roughly 4 % of the overall median).

The impact of temperature on the blank insert TEER values was evaluated. The temperature in the safety cabinet across time was relatively consistent (25.5 °C ± 1.3 °C; values indicate mean and uncertainties one standard deviation); measurements for one time point (Experiment 13 day 17) were performed in a different laboratory and were lower (21.1 °C). A statistically significant trend was observed between the biosafety cabinet temperature and blank insert TEER values (p < 0.001), but it had a low R^2^ value of 0.14. The likely cause of this weak correlation is the temperature consistency. When measurements in the biosafety cabinet were made at the beginning and conclusion of a run in Experiments 6 through 10, they were always within 1 °C, indicating minimal temperature fluctuation within a run. Measurements at different locations within the biosafety cabinet were similarly consistent.

### Biculture samples

The biculture samples maintained consistent average TEER values from 14 d through 22 d (Figure S4). This contrasts with the patterns observed for the monoculture and triculture data that showed different trends during the 22 d period (data not shown). The consistency of the biculture data is why KCl and blank insert data were compared with only the biculture data to evaluate potential correlations.

The biculture data in experiments 1, 2, and 4, taken 1 h or 4 h after the initial measurements, were compared to the values in the first run to assess the within-day variability. The median ratio of the initial to the later mean values was 1.04 (n=17; distribution-free 95 % confidence interval 0.95 to 1.11, treating the 17 comparisons as independent, although they come from three experiments). Ten of the 17 comparisons (59 %) were within 10 % of each other, and 14 (82 %) were within 20 %. The other three ratios were 1.3, 1.5, and 2.

Systematic drift within runs is possible: samples measured later in the run could have consistently higher or lower values than those measured earlier. However, comparing the biculture data across experiments revealed that there was no systematic trend based on the order of the insert analysis (Figure S5).

One possible control measurement to monitor within-run changes in the instrument sensitivity would be to test a selected number of biological samples at the beginning and end of a run. However, when biculture samples were left outside of the incubator for several hours, and measured after each hour, the TEER values decreased with increasing time (data not shown), indicating that this type of control measurement is infeasible because the samples change.

### Correlational analysis

Correlations were not observed between the deviations in the KCl solutions and biculture results compared to their long-term medians (Figure S6). Similarly, comparing deviations in the fitting parameters from the KCl solutions using the Kohlrausch equation (Bockris and Reddy 1970) were also not correlated with deviations in the biculture data (data not shown).

In contrast, the deviations in the blank inserts and biculture data were statistically correlated (p = 7.1 × 10^-6^) (Figure 8). However, the fraction of the total variability that could be explained by the correlation was fairly small (R^2^ = 0.20). When the analysis was performed using only the blank inserts on the plate that had the biculture samples, the correlation was similar (R^2^ = 0.22, p = 5.9 × 10^-6^). This fraction of the total variability was similar to the maximum (R^2^ = 0.23 when comparing monoculture to triculture samples) observed when comparing samples from different gut models tested in the same run (Figure S7). However, comparing biculture samples to monoculture or triculture yielded substantially smaller R^2^ values of 0.06 to 0.10, respectively. When comparing the mean biculture data in a given run to the biosafety cabinet temperature, the correlation was statistically significant but weak (p = 0.04, R^2^ = 0.06).

**Figure 8:**
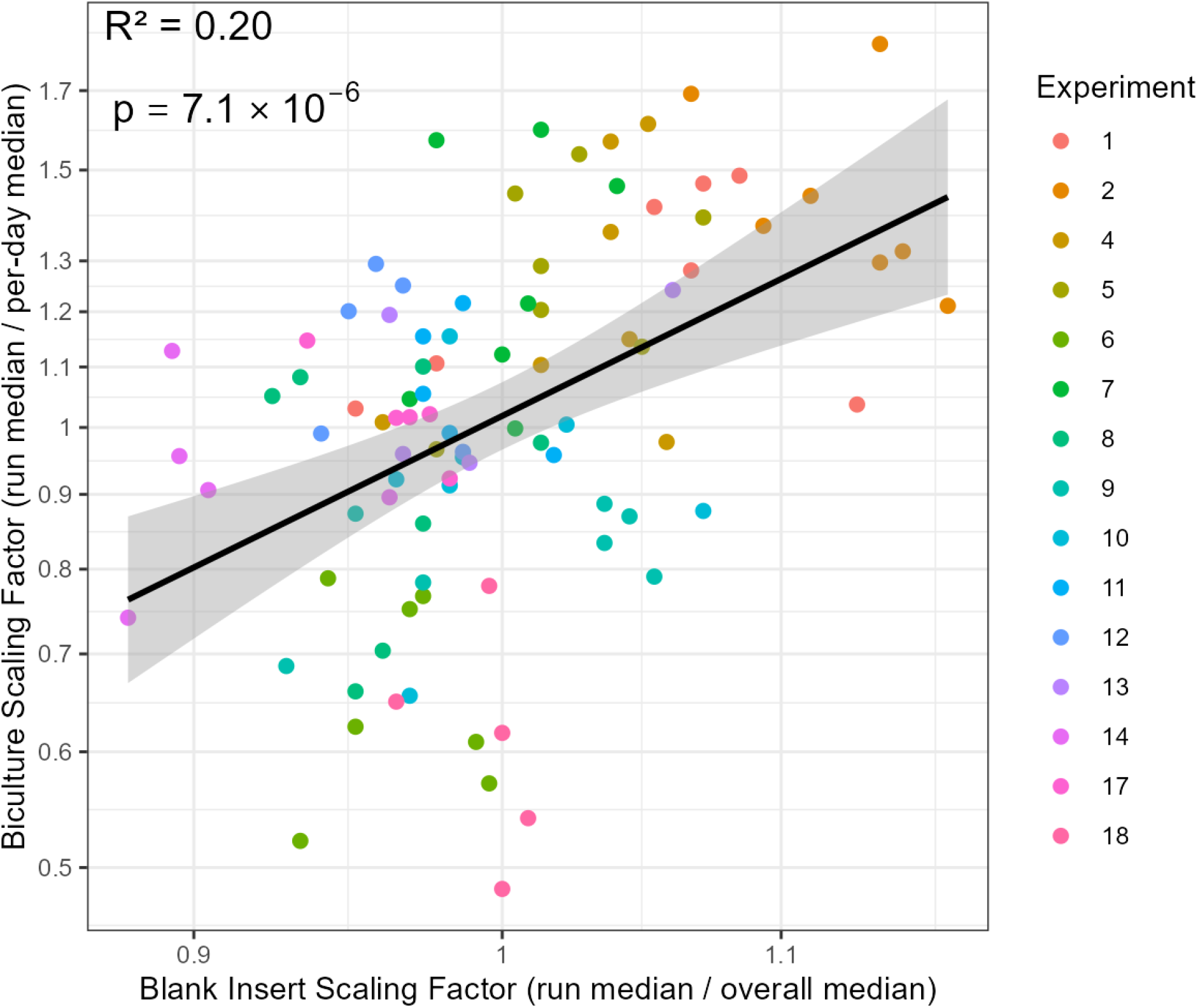
Comparison of run-level scaling factors for biculture cell data and blank insert data. Each point represents one experiment×date combination, colored by experiment (n=92). The x-axis is the median blank insert TEER value for that run divided by the overall median blank value across all runs. The y-axis is the median biculture TEER value for that run divided by the median biculture value for that timepoint across experiments. The black line is the linear regression fit (R² = 0.20, p = 7.1 × 10⁻⁶), indicating a statistically significant but weak association between blank insert and biculture scaling factors. The grey band is the 95 % confidence interval of the fit, and both axes are on a log scale.

## Discussion

Multiple lines of evidence suggest minimal systematic within-run drift: within-run instrument drift evaluations, evaluation of insert order within a plate for the biculture data, and the low plate-to-plate uncertainty contribution for blank inserts. Nevertheless, it could be helpful to analyze the samples in a randomized order, or, alternatively, to vary the order in which samples from different conditions are tested (e.g., test one sample from each treatment, then the second sample from each treatment, etc.) to minimize the effects of unexpected within-run instrument drift. However, if cellular exposure to different concentrations of a test chemical is being evaluated, this may not be advisable due to the potential for the test chemical to transfer among the different inserts, changing the exposure dose. In this case, testing from the lowest to highest concentration may be a better approach.

Multiple analyses were performed to evaluate the potential for the KCl solution and blank-insert control data to correct for run-to-run variability in the biculture data. This was necessary because there was sometimes significant variability in the biological samples among runs within an experiment, as described above for the measurements made 1 h or 4 h apart on the same day. One potential contributing factor to this variability is changes in instrument sensitivity.

One surprising finding was the lack of a correlation between the KCl control measurements and the biculture samples. However, the blank insert values were correlated with the biculture samples (R^2^=0.20). Assuming that the blank inserts are stable for the duration of an experiment, changes in the instrument sensitivity caused by changes in the electrode surface or small variations in the room temperature should have a similar impact on both the blank inserts and biological samples. Variations in the TEER values of biological samples among runs within an experiment could also be due to instability of the biological samples. Some of these potential sources of variation could be minimized by maintaining a consistent schedule of when the medium is exchanged relative to when the TEER measurements are taken, as was done in this study. However, there were no systematic trends in the biculture TEER values between measurements taken prior to or several hours after medium exchange.

Another unexpected finding was that the lowest KCl concentration solutions had more unstable resistance values across a 60 d period compared to higher concentrations (Figure S2). The largest slopes, in terms of their absolute values, were most consistently observed for the 1.25 mmol/L concentration. Chlorine ion autooxidation under typical lab room-temperature and pressure conditions is expected to be extremely slow unless the electrode is positively biased. With a low-ionic-strength electrolyte, slow Ag ion leakage is possible. Evaporation could gradually increase the KCl concentration in the solutions, which would also potentially decrease the solution resistance. Decreasing resistance over time was observed in some samples in a lot, but not in others.

Given their greater instability over time, one recommendation is to avoid using KCl control measurements at the lower concentrations. The instrument manufacturer recommends using higher KCl concentrations of 20 mmol/L and 80 mmol/L (World Precision Instruments 2025). Another recommendation is to use freshly made KCl solutions. When five KCl lots were made at the same time, the coefficient of variation was, on average, 2.9 % among the different concentrations measured the first day they were prepared, a value similar to the lot-to-lot variability values in Figure 4.

However, the KCl solution control measurements were relatively uninformative during the conditions tested in this study. Such measurements could indicate that the electrode needs to be replaced, but that was not needed during this study. It is unclear whether testing multiple KCl solution concentrations provides additional information compared to a single concentration. Reducing the number of KCl concentrations could simplify and shorten the protocol.

Testing the voltohmmeter was similarly uninformative given its stability. This control measurement could be tested periodically or as a follow-up control measurement to help with troubleshooting if an issue is revealed by another control measurement.

Statistical evaluation of biological sample results among different conditions within a run is straightforward, given the limited instrument drift observed. However, the situation is more complicated for comparing results among runs. The run-to-run variability for the blank insert is substantial at approximately 4 % of the median value. This value is probably an underestimate of the true magnitude of the run-to-run variability since it is confounded with other types of noise. Additional contributing factors could contribute to run-to-run variability of biculture samples, such as random changes to the biological sample barrier integrity.

A simple statistical test, such as a t-test, would be inappropriate for comparing biological sample results among different runs, because it would not account for the run-to-run variability of the TEER measurements. There are several paths to make such comparisons. One possibility is to compare the magnitude of the effect size to estimates of the run-to-run variability. If the effect size is substantially larger than the run-to-run variability (estimated by prior information), the results are more likely to be significantly different. Another possibility is to combine results from multiple runs, potentially including samples measured on different runs on the same day, since the random effects from run-to-run variability can be accounted for using analysis of variance (ANOVA).

### Future work

One valuable topic for future work is to further investigate the potential to use blank inserts to better control for run-to-run variability for the biological samples. The intended purpose of the inserts during this study was solely for background subtraction during an experiment. This is why the inserts were kept in DMEM complete medium, despite the potential for membrane pore clogging, and were replaced each experiment. Determining the long-term stability of a set of inserts stored between runs in a simpler medium less likely to foul the inserts, such as DMEM without FBS, could be beneficial to correct for run-to-run variability caused by changes in electrode sensitivity.

Another related possibility is the development of a control material with demonstrated homogeneity and stability. While the CaliCell insert can be used with chamber systems, there are not similar materials to our knowledge for probe-based systems and only a single resistance is available for the chamber system. This material should be stable across a long time period, enabling its use across multiple experiments. It could potentially have a higher resistance value than those measured in this study for the blank inserts, which were substantially smaller than those for the biculture samples. The higher relative uncertainty contribution for repeated measurements of the blank inserts compared to repeated measurements of KCl concentrations could have stemmed from the lower blank insert TEER values that were closer to the detection limit. While heterogeneity in the insert could contribute to repeated-measurement uncertainty, placing inserts in the chamber in a consistent orientation, as was done in this study, should minimize this contribution.

One substantive challenge with making such a control material is that TEER measurements are made on inserts varying significantly in size from 6-well plates to 96-well plates. It is unclear if a single control material could be used across inserts with different sizes and probe types.

While temperature is well known to impact TEER measurements (Blume et al. 2010; Sharma et al. 2025), the relatively consistent temperatures in this study resulted in weak correlations between temperature and the blank insert or biculture TEER values. This suggests that temperature fluctuations played a relatively minor role in the run-to-run variability in this study. Nevertheless, other laboratories may have significantly larger seasonal temperature fluctuations. There are two potential strategies to reduce temperature fluctuations: 1) use a hot plate set at 37 °C to maintain the temperature of the cells at the same temperature as the incubator (Vincentini et al. 2022; De Angelis et al. 2023), or 2) use an automated TEER system that fits within a temperature-controlled incubator. One limitation of using a heating plate is that this is not possible for chamber TEER systems. Neither of these options were investigated in this study but could be explored in future work. Another possibility is to correct for the temperature-induced bias of TEER measurements (Blume et al. 2010).

Overall, this study provides a case study for evaluating the sources of variability in a single TEER system. Several approaches for assessing within-run and run-to-run variabilities have been described in this study. The method used to evaluate instrument drift within and among runs using a stable test material or solution could be used to evaluate other TEER systems. The number of measurements needed to sufficiently characterize the within-run and run-to-run variability would need to be assessed on a case-by-case basis.

Several potential sources of variability were found in this study to be relatively minor: voltohmmeter, temperature, and within-run variability. Run-to-run variability was consistently substantial and was not reduced by more frequent repolarization or reconditioning of the electrodes. Run-to-run variability of the biological samples could be partially explained by the blank-insert measurements. A different control-measurement strategy, such as using control materials or inserts stored in a simpler medium, could further clarify the contribution of measurement variability caused by changes in electrode sensitivity. This is particularly important given that the primary strategy to characterize changes in electrode sensitivity attempted here, measurements of KCl solutions, was unsuccessful because these results did not correlate with the biculture data.

Developing documentary standards for characterizing TEER systems could be a valuable step toward improving the reproducibility of these measurements and reducing laboratory-to-laboratory variability. In an interlaboratory study evaluating air-liquid interface models, the values for the control cells were relatively consistent within laboratories but differed by a factor of approximately 3 among laboratories (Braakhuis et al. 2023). Metrics to characterize the within-run and run-to-run variability could be a useful addition to TEER reporting standards (Sharma et al. 2025).

## Declarations

## Acknowledgements

We sincerely thank NIST colleagues John Elliott, Anthony Asmar, and Ana Barrios for helpful discussions regarding this project, valuable input on a manuscript draft, and for performing preliminary experiments, respectively.

## Disclaimer

The views in this publication are those of the authors and not an official position of the United States Government. The authors declare no competing financial interest. Certain commercial equipment, instruments, or materials are identified in this paper in order to specify the experimental procedure adequately. Such identification is not intended to imply recommendation or endorsement by the National Institute of Standards and Technology, nor is it intended to imply that the materials or equipment identified are necessarily the best available for the purpose.

## Ethics approval

The National Institute of Standards and Technology Research Protections Office determined this project is ‘not human subjects research’ (Protocol No. MML-2021-0296).

## Data availability

The data analyzed in this study is available in the Supporting Information.

## Supporting figures

**Figure S1.**
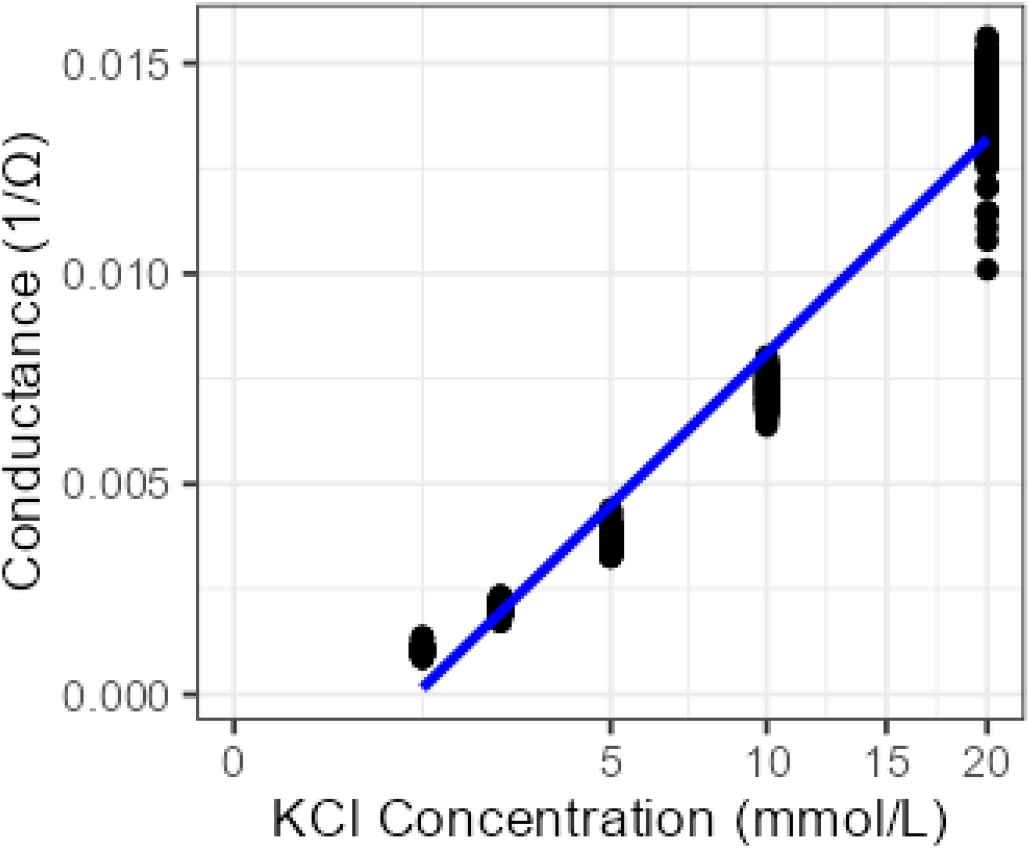
Plot of the KCl concentrations (x-axis is on a square-root scale) versus conductance. The blue line depicts the best-fit linear model for Kohlrausch’s law. Each dot is a single measurement; three replicate measurements were made of each KCl solution in each run. Data are from 22 experiments (21 at 1.25 mmol/L), with 396 to 429 measurements per concentration.

**Figure S2.**
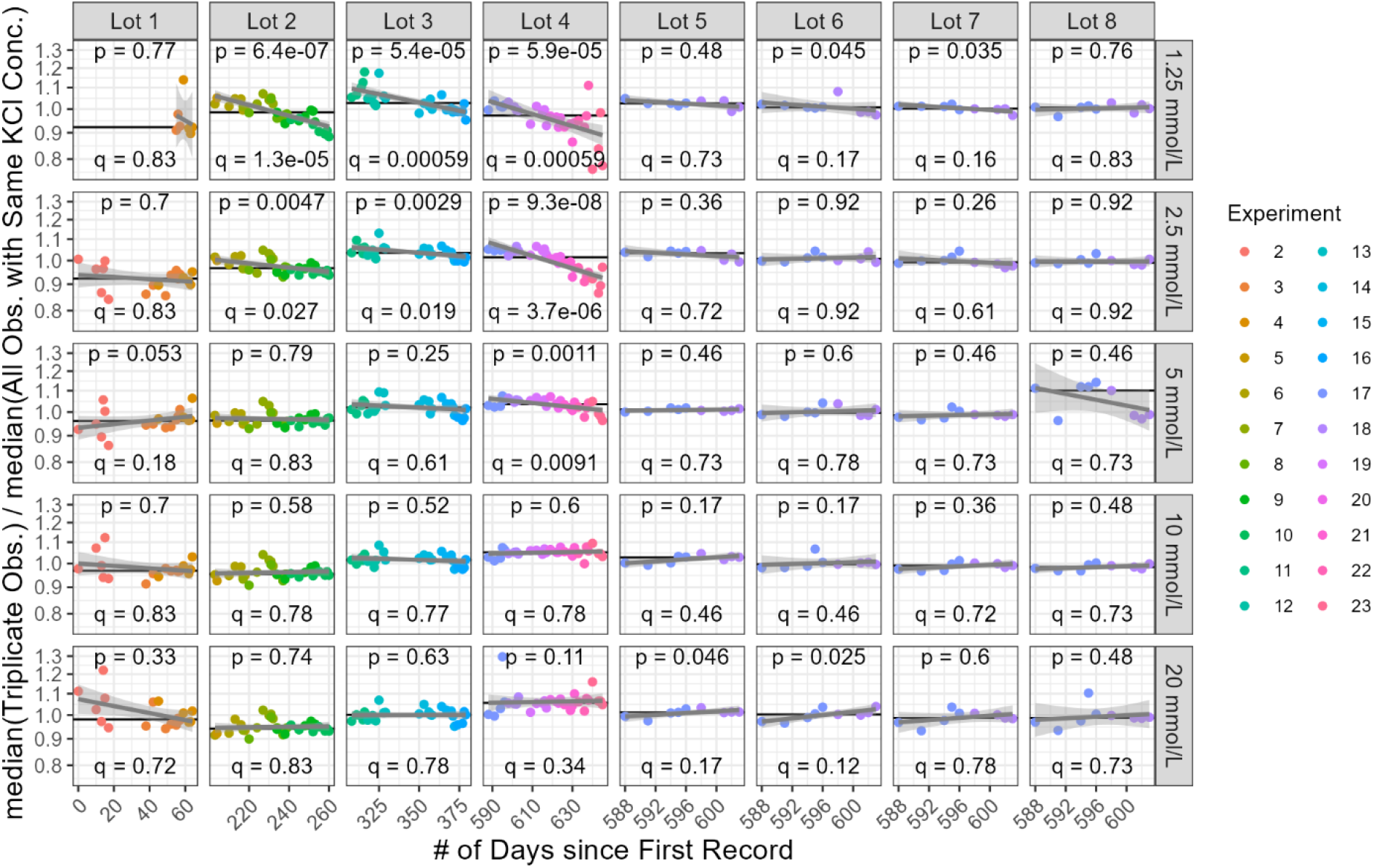
Evaluation of temporal trends in KCl data for different lots across 22 experiments. Each panel represents one lot×concentration combination (n=3 measurements for a single solution for each run). The color of the data points indicates the experiment. The x-axis is the number of days since the first experiment; the y-axis is the median of triplicate measurements divided by the overall median for that KCl concentration. Grey lines and bands show linear regression fits and their 95 % confidence intervals. The y-axis is on a log scale, and the black horizontal line is the median for that lot and concentration divided by the overall median for that concentration. Each panel reports the p-value from Kendall’s rank correlation test and the corresponding q-value, which estimates the expected proportion of false positives among all tests called significant at that threshold.(Storey 2003)

**Figure S3.**
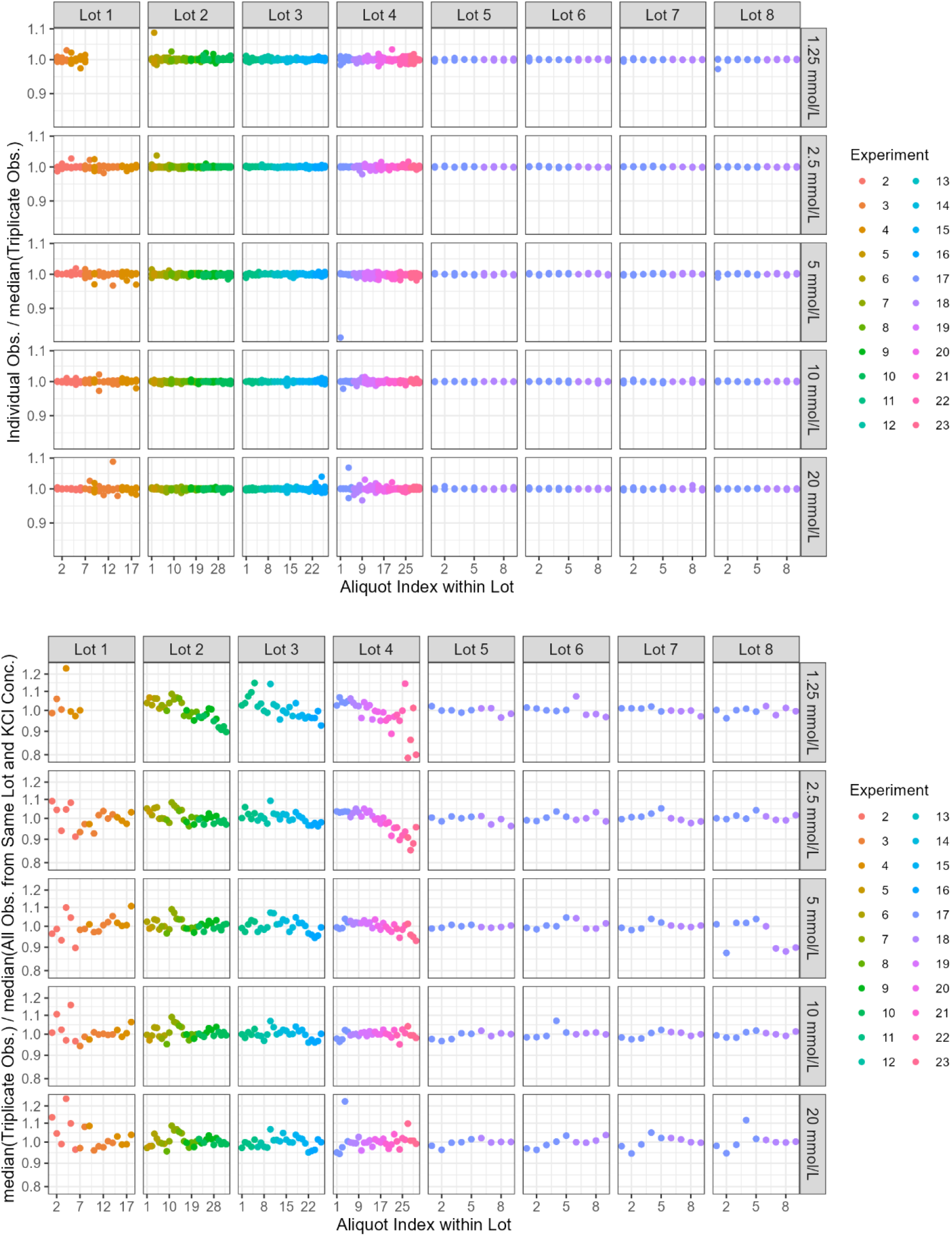
Control charting of KCl measurements organized by experiment, KCl concentration (mmol/L), and the KCl lot (1 through 8). The top panel plots each individual measurement divided by the median of its triplicate set, showing within-triplicate variability. The bottom panel plots the median of each triplicate set divided by the overall median for that lot and concentration, showing aliquot-to-aliquot variability. Both y-axes are on a log scale. Within each lot, aliquots are numbered in order of measurement date.

**Figure S4.**
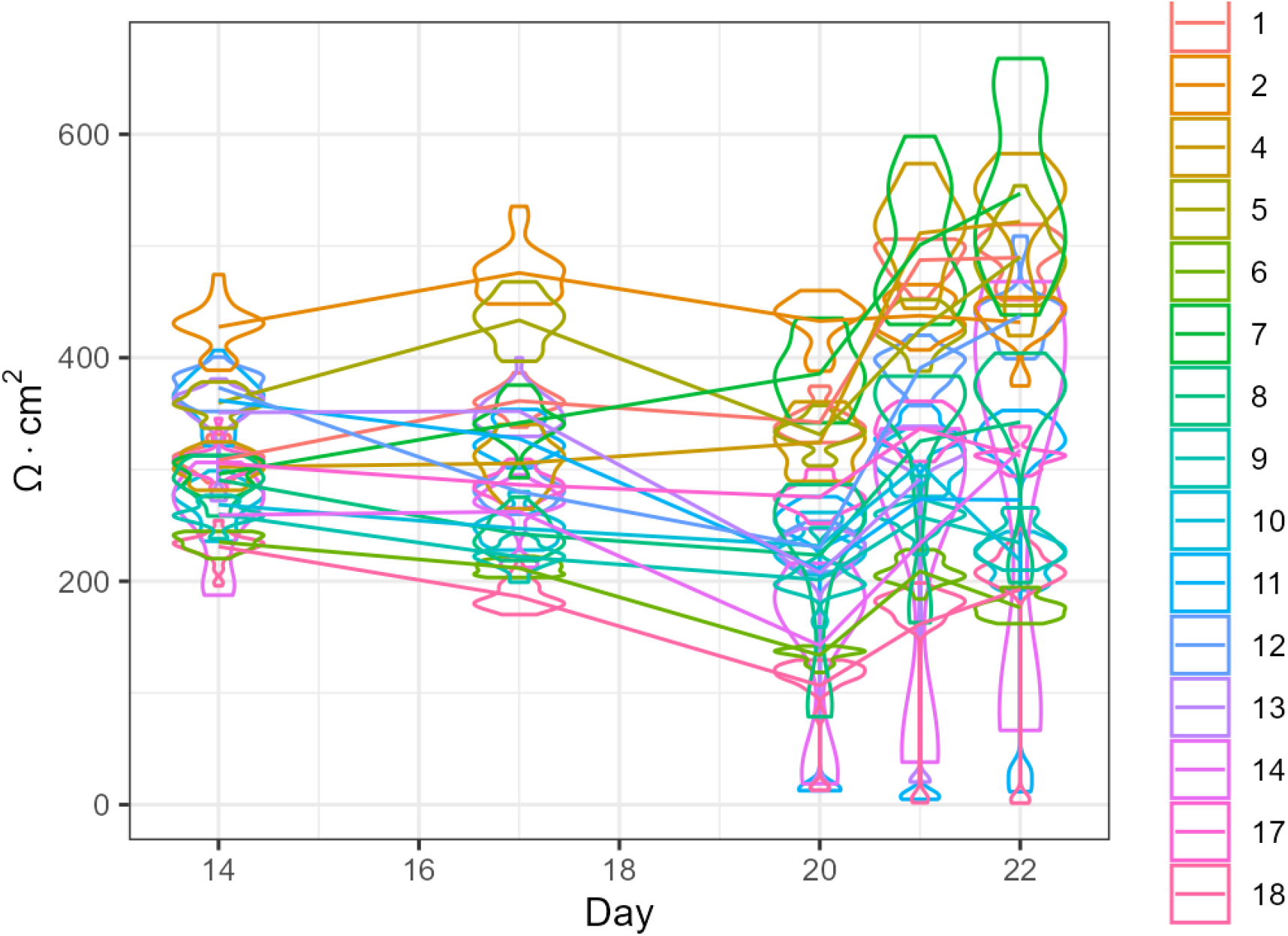
Violin plots of biculture TEER measurements at 14 d, 17 d, 20 d, 21 d, and 22 d. Each violin represents the distribution of measurements for one experiment (n=3 measurements for each of 9 to 22 inserts per run) at one timepoint, colored by experiment. Lines connect each experiment’s mean across timepoints.

**Figure S5.**
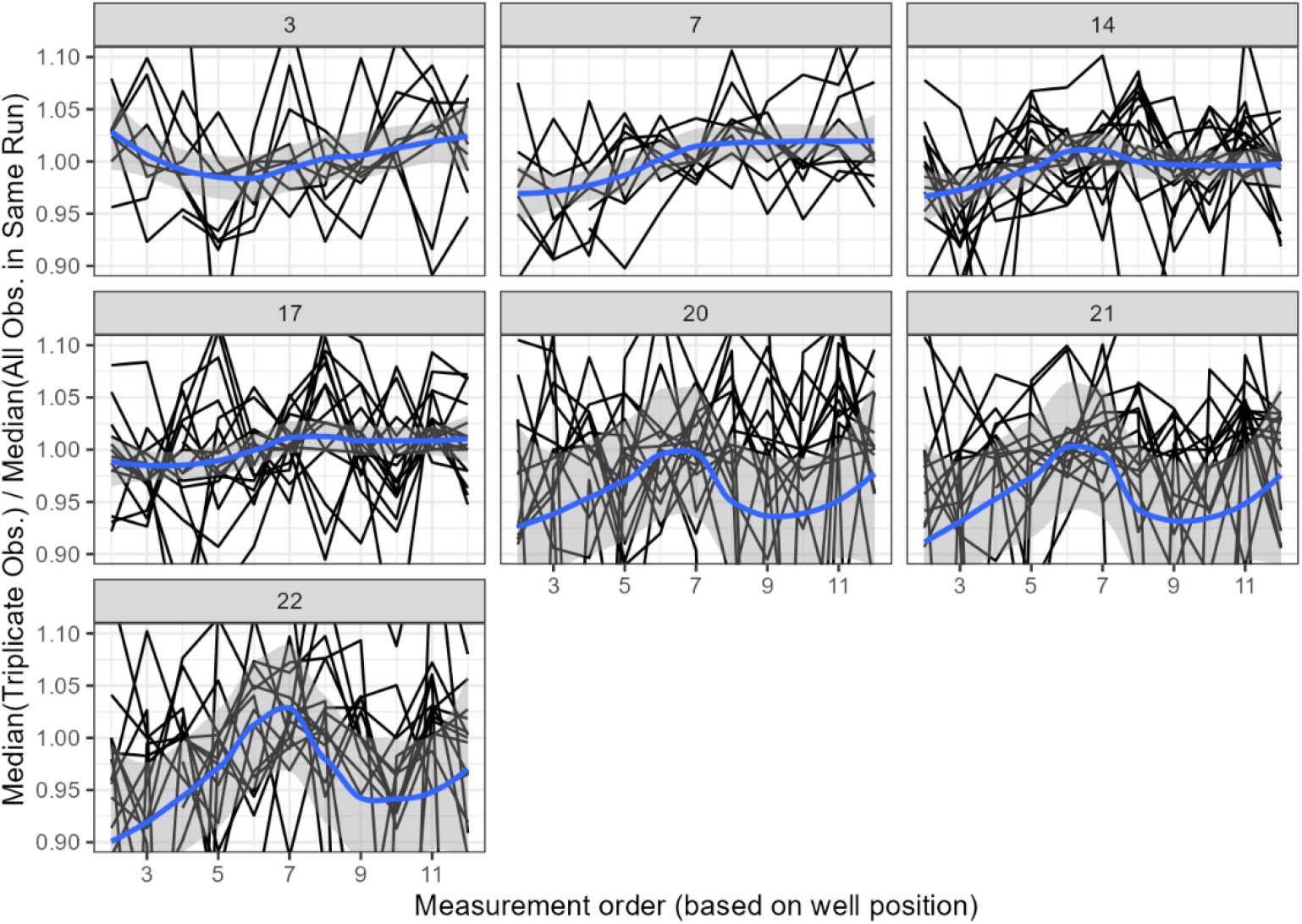
Evaluation of the impact of measurement order on the biculture cell model TEER values for different time points: 3 d, 7 d, 14 d, 17 d, 20 d, 21 d, and 22 d. The x-axis shows the measurement order based on well position. The y-axis shows the median of triplicate measurements for a biculture sample divided by the median value for that run. Each black trace represents a different experiment×day×plate combination. The blue line is a loess smoothing function, and the grey band is its 95 % confidence interval. The y-axis is limited to 0.90 to 1.10; 11 % of values lie outside this range.

**Figure S6.**
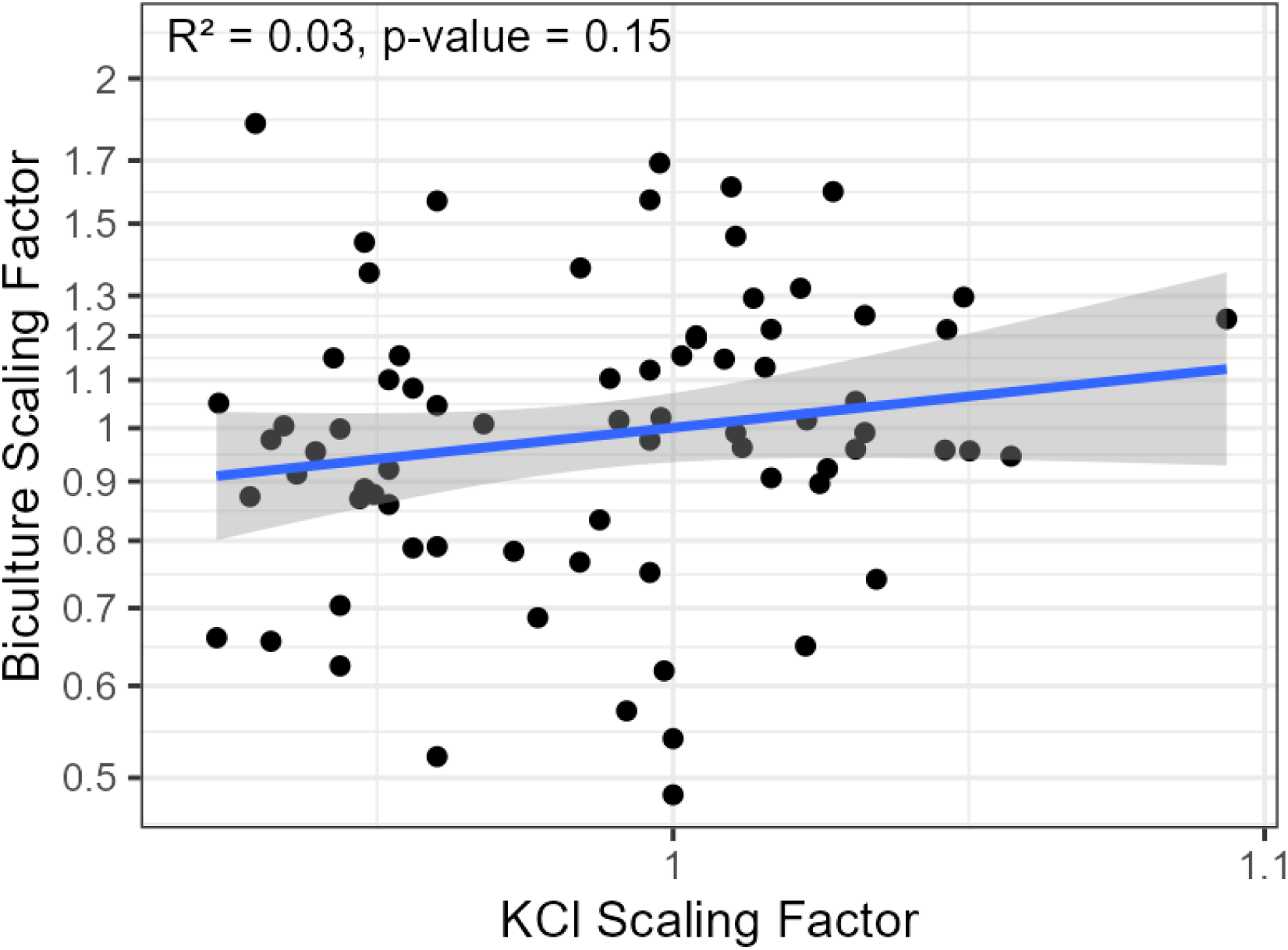
Comparison of run-level scaling factors for KCl and biculture data. Each point represents one experiment×day of experiment (e.g., 14 d after the experiment started) combination (n=75). The x-axis is the median, over all KCl measurements in that run (all concentrations), of each resistance divided by the overall median for its concentration. The y-axis is the median biculture TEER value for that run divided by the median biculture value for that timepoint across experiments. Both axes are on a log scale. The blue line is the linear regression fit and the grey band is the 95 % confidence interval (R^2^ = 0.03, p-value = 0.15), indicating no significant association between KCl and biculture scaling factors.

**Figure S7.**
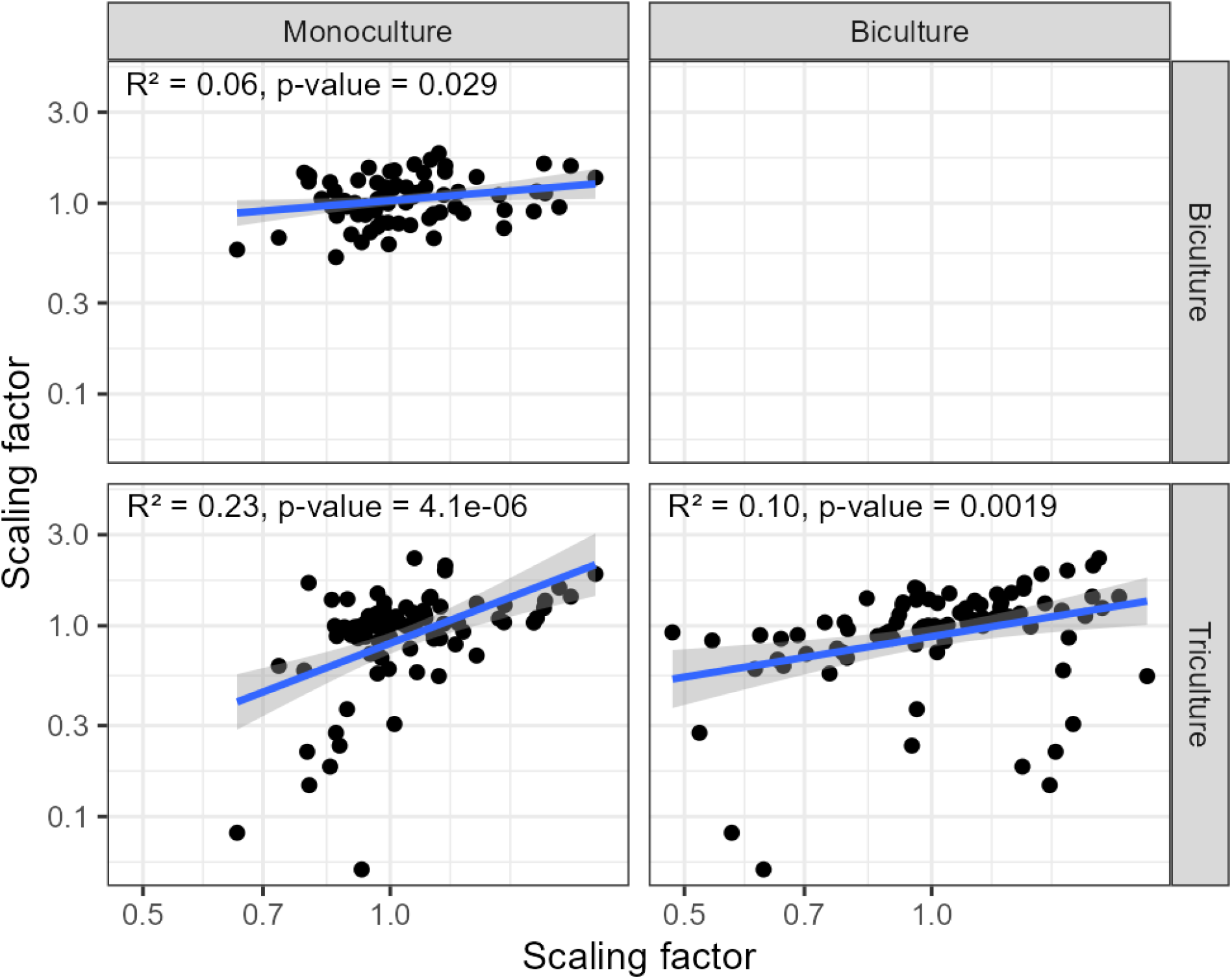
Pairwise comparisons of run-level scaling factors for monoculture, biculture, and triculture data. Each scaling factor is the median TEER value for that experiment×day of experiment (e.g., 14 d after experiment started) combination divided by the median for that cell type and timepoint across all experiments (n=77 to 91 per panel). Each point represents one experiment×day combination. The blue line is the linear regression fit and the grey band is the 95 % confidence interval. R^2^ values and p-values are shown in each panel. Both axes are shown on a log scale.

